# *In silico* investigation of *Aedes aegypti* male-determining factor (NIX): RNA recognition motif-3, structural model and selective nucleic acid binding mode

**DOI:** 10.1101/2020.11.13.381210

**Authors:** Monika A. Coronado, Danilo S. Olivier, Kelly C. Borsatto, Marcos S. Amaral, Raghuvir K. Arni, Raphael J. Eberle

## Abstract

Mosquito borne viruses and their corresponding human infections have a fast growing impact on worldwide public health systems; in particular, *Aedes aegypti* and *Aedes albopictus* are transmitting these viruses. Only female mosquitoes bite and spread diseases. Male development in *A. aegypti* is initiated by male determining factors. These factors are potential candidates for the implementation of vector control strategies. Where at an early stage in gender development female mosquitoes are converted into harmless males. Among these factors, a novel gene, NIX, has been identified that shares moderate identity with transformer-2, one of the key sex-determination genes in *Drosophila melanogaster*. Hall et al. 2015 described two RNA recognition motifs (RRMs) in the NIX sequence, but a UniProt database search showed that *A. aegypti* NIX contains three RNA RRM. A homology model of NIX_RRM-3 was generated to investigate the interaction with RNA using molecular dynamics simulations. The sequence AGACGU was the most interesting result. During the binding process A1, G2 and G5 adopt an unusual *syn* conformation. The results let assume that the C-terminus of NIX_RRM-3 is involved in the recognition process of the target RNA motif (e.g. AG) and controls the interaction between target RNA and *A. aegypti* NIX.

## 1. Introduction

Climate change has a profound effect on the global occurrence and burden of infectious diseases [1–3], especially mosquito borne viruses. The resulting altered distribution of viruses such as Dengue, Yellow Fever, West Nile, Chikungunya, Zika and Japanese Encephalitis and their vectors (e.g. *Aedes aegypti* and *Aedes albopictus*) are key factors of public health that can dramatically change infection patterns and require attention [4–6].

The recent strategies to control mosquitoes that transmit viruses are based on genetic techniques such as the sustained release of sterile or transgenic “self-limiting” mosquitoes [7]. Given the fact that only female mosquitoes bite and spread diseases, there has been substantial interest in manipulating mosquito sex determination using these genetic techniques and others, including gene drive [8,9]. Therefore, elucidating the genetic basis for sex determination could, for instance, facilitate the male-only production of cohorts for release, or allow transformation of mosquitoes with sex-specific “self-limiting” gene cassettes [10].

Male development in *Aedes aegypti* (*A. aegypti*) is initiated by male factors (M-factor) located on the homomorphic sex-determining chromosome within a Y chromosome-like region referred to as the M-locus [11,12]. Consequently, a mosquito M-factor would be useful in implementing vector control strategies where female mosquitoes at an early stage in gender development are converted into harmless males [13]. A number of male-specific genomic sequences were identified in *A. aegypti* [14]. Among these, a novel gene classified as Nix that shares moderate identity with transformer-2, one of the key sexdetermination genes in *D. melanogaster* [15]. The *A. aegypti* Nix gene is 985 bp in length and encodes a polypeptide with 288 amino acids including two RNA recognition motifs (RRMs). Nix exhibits persistent male linkage and is expressed early in embryonic development [14], being essential characteristics of the M-factor. Furthermore, somatic knockout of Nix in *A. aegypti* male embryos results in feminization whereas ectopic expression of Nix in female embryos leads to masculinization, clearly demonstrating that Nix is instrumental for the initiation of male development [14]. Nix is hypothesized to be a splicing factor that acts directly or indirectly on the doublesex (dsx) and fruitless (fru) genes involved in sex-determination [16].

So far, no structural information is available about the *A. aegypti* Nix protein and its interaction with the target RNA. The investigation of the binding specificity between protein and nucleic acids can help to understand the interaction process and conformational changes in the protein and the RNA, respectively.

Analyses of the UniProt database (https://www.uniprot.org/) demonstrates that the *A. aegypti* NIX protein sequence contains an additional RRM to the two RRMs described previously [14]. A model of the full-length protein and the three RRM domains were constructed. Molecular dynamics (MD) simulations were performed with the NIX_RRM-3 domain to observe the stability of the model over time. The NIX_RRM-3 model was used for docking and MD simulations with a set of various RNA sequences with the goal of identifying an appropriate target RNA sequence and exploring the binding process between the protein and the RNA. AGACGU showed a robust interaction with NIX RRM-3. Interestingly A1, G2 and G5 bases undergo an unusual *syn* conformation during the interaction with the C-terminus of NIX_RRM-3. We suppose that the NIX_RRM-3 domain in NIX is essential for identification and binding of the target RNA.

## 2. Material and Methods

### 2.1 *In silico* analysis

Multiple sequences of NIX, male factor, transformer, and RNA binding proteins were retrieved from the NCBI database and sequence alignments were performed using the MUSCLE [17] and Box Shade (http://www.ch.embnet.org/software/BOX_form.htm) web servers. A homology model of full-length *A. aegypti* NIX (GeneBank: AHW46195.1 Uniprot: A0A0F6MY85) as well as *A. aegypti* NIX_RRM-1, −2, and −3 domains were created using the Zhang lab I-tasser online tool [18].

### 2.2 Docking and molecular dynamics

The HADDOCK2.2 [19] web server was used to perform the molecular docking between the *A. aegypti* NIX_RRM-3 protein domain and the following RNA sequences (AAGAAC, AGACGU and AAACGU). To choose the start configuration, energy values for the protein-RNA interaction as well as visual inspection was used. Molecular dynamics simulations were performed using the Amber18 [20] software package. The descriptions of all-atoms interactions were performed using the FF14SB [21] force field for the protein while the RNA sequences were described using the OL3 [22] force field. The NIX_RRM-3 model was created using the Zhang lab I-tasser online tool [18–19]. The protonation state for the NIX_RRM-3 protein was settled using H++ [23] web server at pH 7.5. The initial protein model was submitted to MD simulations for 300 ns to obtain a relaxed structure and the model quality monitored by PROCHECK [24]. NIX_RRM-3 and the protein/RNA-complexes were neutralized with Na^+^ or Cl^-^ ions and, placed in an octahedral box with TIP3P water extended 10 Å from the protein atoms. Energy minimization was performed in two steps to remove bad contacts. Initially, both the protein and the protein/RNA complex were constrained and a constant force of 50.0 kcal/mol- Å^2^ was performed for 5000 steps of steepest descent succeeded by 5000 steps of conjugate gradient, following which, 10000 steps energy minimization without constraints were conducted. The system was heated from 0 to 298 K under constant atom number, volume and temperature (NVT) ensemble, maintaining the protein and protein/RNA complex restrained with a constant force of 10.0 kcal/mol- Å^2^. Thereafter, an equilibration step was performed under constant atom number, pressure and temperature (NPT) ensemble for 500 ps and the simulation was performed for 200 ns with 2 fs time steps. Constant temperature (298 K) and pressure (1 atm) were controlled by Langevin coupling. Long-range electrostatic interactions were treated via Particle-Mesh Ewald (PME) [25] while a cut-off distance of 10 Å was attributed to Van der Waals interactions.

### 2.3 Molecular dynamics analyses and Energy Interaction calculations

The MD simulations were analyzed using CPPTRAJ [26] of the AmberTools18 package. System equilibration and convergence were assessed by analysis of the root mean square deviations (RMSDs). Protein and RNA flexibilities were studied by root mean square fluctuation (RMSF). RMSFs were calculated residue-by-residue over the equilibrated trajectories. Radius of Gyration (RoG) and surface area were calculated to check major structural changes in the protein. The molecular mechanics/generalized Born surface area (MM/GBSA) energy was calculated between the protein and the RNA molecules considering the stable regime using the last 20 ns of the MD simulations and stripping all the ions and water molecules.

## 3. Results and Discussion

### 3.1 Sequence analysis of full length *A. aegypti* NIX protein and RRM domains 1-3

*A. aegypti* NIX is a 288 amino acid polypeptide that has been reported of comprising two RNA recognition motifs (RRMs). The first RRM is comprised of the region Tyr19 to Ser94 and the second RRM of the region Arg205 to Lys282 [14]. However, a search on the UniProt database (https://www.uniprot.org/), demonstrated that *A. aegypti* NIX, contains a third RRM motif (Ile108 to Arg179). The schematic construction of the NIX protein with RRM-1 to RRM-3 are presented in Fig. 1A and 1B.

**Fig. 1.**
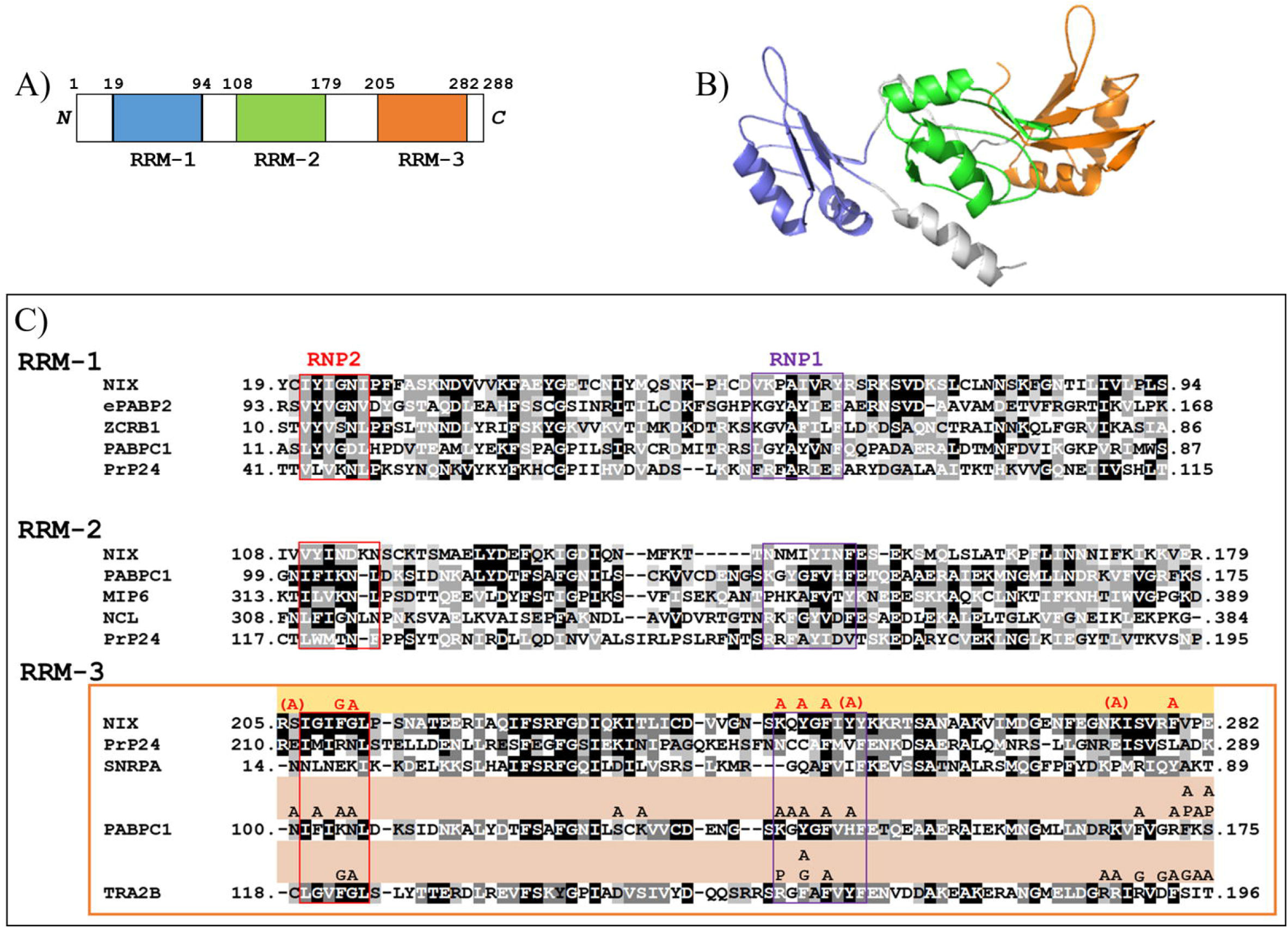
Primary and tertiary structures of *A. aegypti* NIX. (A) Schematic diagram of *A. aegypti* Nix. The three RRM domains are represented in different colors (RRM-1 (blue), RRM-2 (green), and RRM-3 (orange)) and, amino acid residue numbers indicate the approximate boundaries of the three RRMs. (B) Homology model of Nix, the RRM domains; color coded as in A. (C) Sequence alignment of NIX_RRM-1 to RRM-3 with different RRMs. RNP1 and RNP2 motifs are highlighted (purple and red, respectively). The sequence of NIX_RRM-1 is aligned with *Xenopus laevis* Embryonic polyadenylate-binding protein 2-B (ePABP2), RNA-binding motif-containing protein 1 (ZCRB1), *Homo sapiens* Polyadenylate-binding protein 1 (PABPC1), *Saccharomyces cerevisiae* U4/U6 snRNA-associated-splicing factor (PrP24). The NIX_RRM-2 sequence is aligned with *Homo sapiens* Polyadenylate-binding protein 1 (PABPC1), *Saccharomyces cerevisiae* RNA-binding protein (MIP6), *Mesocricetus auratus* Nucleolin (NCL), *Saccharomyces cerevisiae* U4/U6 snRNA-associated-splicing factor (PrP24). The NIX_RRM-3 sequence (orange box) is aligned with *Saccharomyces cerevisiae* U4/U6 snRNA-associated-splicing factor (PrP24), *Homo sapiens* U1 small nuclear ribonucleoprotein A (SNRPA), *Homo sapiens* Polyadenylate-binding protein 1 (PABPC1), *Homo sapiens* Transformer-2 protein homolog beta (TRA2B). The amino acids involved in nucleic acid binding and the corresponding bases from PABPC1 and TRA2B crystal structures (PDB code: 1CVJ, 2RRA and 2KXN) are highlighted, adenine (A), guanine (G) and phosphate backbone (P).

A typical RRM domain consists of 80 to 90 amino acid residues [27]. However, the *A. aegypti* NIX_RRMs are slightly smaller whereas RRM-1 comprises 75 residues, RRM-2 of 71 and RRM-3 of 77 respectively. Nevertheless, all three RRMs domains contain the characteristic RNA nucleoprotein motifs 1 and 2 (RNP1 and RNP2) as indicated in Fig. 1C. The RNP motifs in the RRMs are necessary for their specific RNA-recognition modes [27]. The consensus RNP2 and RNP1 sequences are defined as [I/L/V]-[F/Y]-[I/LV]-X-N-L and [K/R]-G-[F/Y]-[G/A]-[F/Y]-[I/L/V]-X-[F/Y] [27]. Based on the sequence alignment of the three NIX_RRMs and related protein structures we identified the *A. aegypti* NIX_RRM-1 (RNP2 and RNP1) motifs sequences are I-Y-I-G-N-I and V-K-P-A-I-V-R-Y, respectively. NIX_RRM-2 presents the motif sequences of V-Y-I-N-D-K and N-N-M-I-Y-I-N-F for RNP2 and RNP1, respectively. The sequences for NIX_RRM-3 (RNP2 and RNP1) were identified as I-G-I-F-G-L and K-Q-Y-G-F-I-Y-Y, respectively (Fig. 1C). The *A. aegypti* NIX RNP2s and RNP1s differs from the typical consensus sequences described by Muto and Yokoyama (2012) [27]. NIX RNP2 and RNP1 sequences of RRM-3 domain showed a high similarity with the corresponding RNPs of *Homo sapiens* Transformer-2 protein homolog beta (TRA2B), with 3D structures in complex with RNA (PDB codes: 2RRA and 2KXN).

### 3.2 *A. aegypti* NIX_RRM-3 model generation, molecular dynamics simulation and validation of the predicted structure

The NIX_RRM-3 model was generated using the Zhang lab I-tasser online tool [18]. Molecular dynamics (MD) simulations were performed to investigate the structural and dynamical changes over time in the NIX_RRM-3 model. The equilibration of the simulations was monitored by calculating the root mean square deviation (RMSD) and the root mean square fluctuation (RMSF) for the backbone atoms with respect to the initial conformation over time with the aim of verifying that all simulations maintained confirmations that were stable during an 300 ns simulation (Supplementary Fig. S1). The NIX_RRM-3 model after MD simulations is presented in Fig. 2A.

**Fig. 2.**
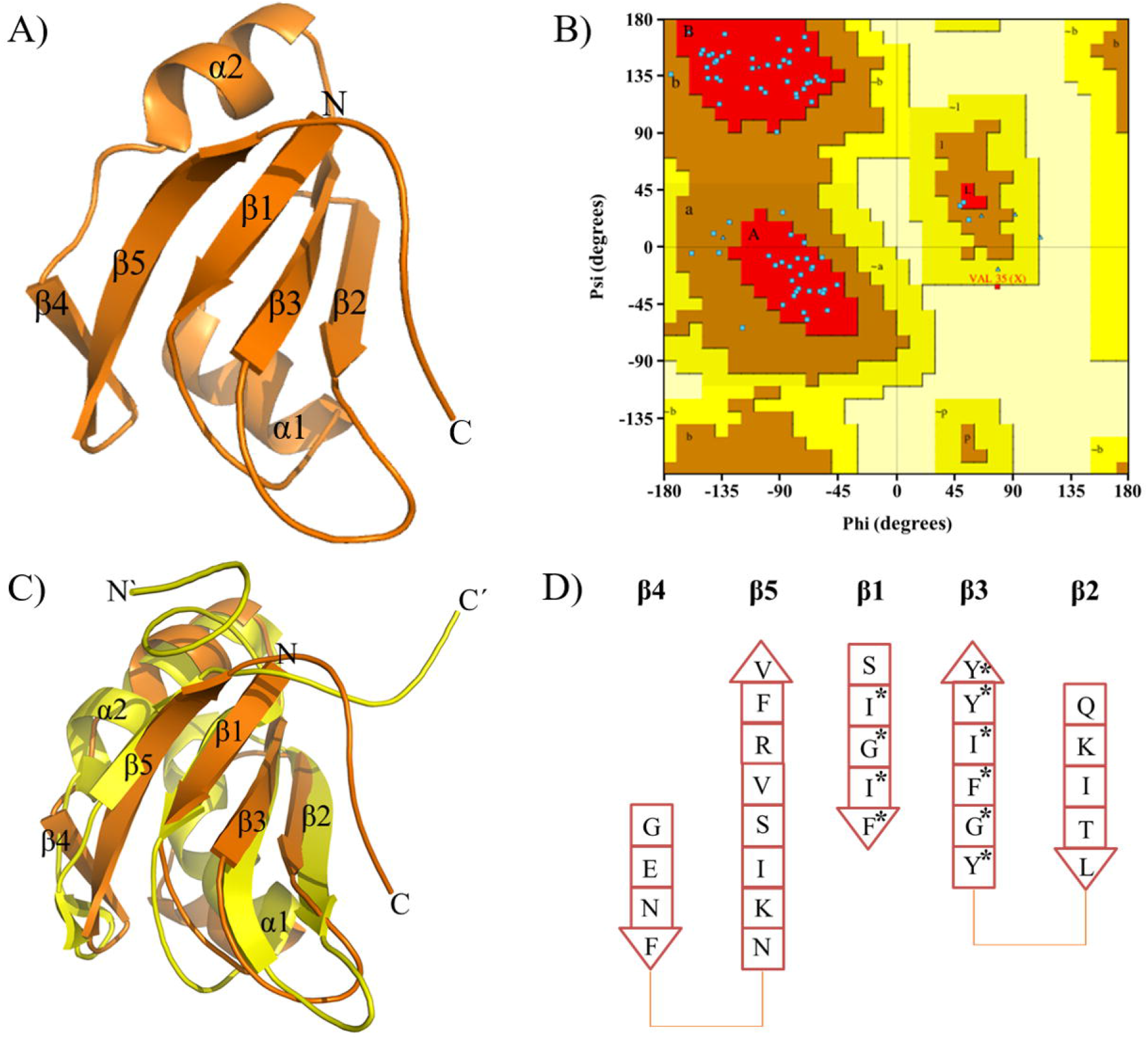
3D model of *A. aegypti* NIX_RRM-3. (A) Homology model after MD simulations of NIX_RRM-3 domain. (B) Ramachandran plot of NIX_RRM-3 after MD simulations. (C) Structural overlay of NIX_RRM-3 (orange) with human TRA2B (PDB entry: 2KXN) (yellow). (D) Scheme of the NIX_RRM-3 β-sheet distribution, some of the amino acids which form the RNP2 (β1) and RNP2 (β3) are highlighted (asterisk).

The evaluation of the stereochemical properties of the predicted *A. aegypti* NIX_RRM-3 model was performed using PROCHECK [24]. The corresponding Ramachandran plot demonstrates that the conformations of 90 % of the residues are located in the favored regions, 9.0 % are in the allowed regions and only 1.0 % were flagged as outliers (Fig. 2B). The stereochemical parameters for the omega angle is −1.3, bad contact are −0.4 and the overall G-factor is 0.3. The final model consists of 83 residues. The structural overlay of the NIX_RRM-3 model and the related structure of the human TRA2B (PDB entry: 2KXN) (RMSD 2.85 Å) is presented in Fig. 2C and supplementary Fig. 2.

The canonical RRM structure consists of a four-stranded antiparallel β-sheet packed against two α-helices with a typical βαββαβ topology [27] whereas the four-stranded β-sheet is the primary RNA-binding surface [28]. The modeled domains of NIX_RRM-1 and −3 contain an additional β-strand, which is located in the loop between α-helix 2 and the C-terminal β-strand. NIX_RRM-2 possesses the typical topology of the RRM family (Supplementary Fig. S3 and Fig. 1B). The human TRA2B and Polyadenylate-binding protein 1 (PABPC1) protein structures contain also an additional β-strand at the same position as the NIX_RRM-3 model [29,30] (Fig. 2C and supplementary Fig. S2). Human TRA2B and PABPC1 structures in complex with RNA showed that the nucleic acids mainly interact with the core βαββαβ and the additional β-strand seems to have a stabilizing function for the RNA interaction [29,30] (Supplementary Fig. S4). The core of the βαββαβ topology is formed by β1 and β3; RNP2 and RNP1 are located in these β-strands, which are also shown for NIX_RRM-3 (Fig. 2C and D).

### 3.3 Identification of a target RNA sequence interacting with the *A. aegypti* NIX_RRM-3 model

A computational approach was used a) to identify a target RNA sequence that interacts with NIX_RRM-3, and b) to study their corresponding interactions. For this, a RNA sequence was inserted near the RNP1 and RNP2 motifs of NIX_RRM-3 using molecular docking. Subsequent MD simulations were carried out to provide statistical results to better understand the interactions between the RRM-3 domain and the RNA. It has been suggested that the RNA sequence AAGAAC is utilized to interact with human TRA2B (PDB entry: 2KXN). Based on the similarities between NIX_RRM-3 and TRA2B RNPs, the RNA sequence was used for a primary MD simulation with NIX_RRM-3. The analysis of the RMSD and RMSF showed a strong fluctuation during 200 ns of MD simulations (Supplementary Fig. S5) and the RNA molecule shifted >20 Å away from the starting position (Table 1 and supplementary Fig. S9). Based on those primary results the RNA sequence AGACGU was chosen for further MD simulations with NIX_RRM-3. The human TRA2B recognize mainly AGAA of AAGAAC [29] and Auweter et al. (2006) suggested a default RNA binding sequence for an RRM that might be a dinucleotide A/C-G in N1-N2 position [28]. The first three bases of AGACGU are similar to the AGAA sequence recognized by human TRA2B and the last three bases contain the CG pattern postulated by Auweter et al. (2006). Interestingly the RMSF fluctuation of the NIX_RRM-3/AGACGU complex reduced dramatically (Supplementary Fig. S6) compared with the NIX_RRM-3/AAGAAC complex (Supplementary Fig. S5).

**Table 1.**
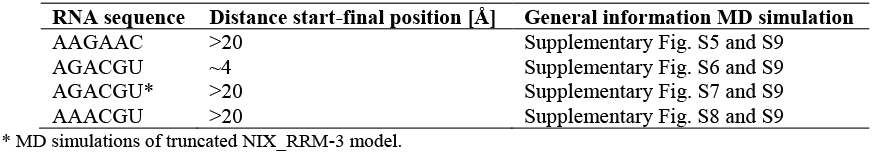
RNA sequences used for MD simulations with *A. aegypti* NIX_RRM-3 model.

| RNA sequence | Distance start-final position [Å] | General information MD simulation |
| --- | --- | --- |
| AAGAAC | >20 | Supplementary Fig. S5 and S9 |
| AGACGU | ~4 | Supplementary Fig. S6 and S9 |
| AGACGU* | >20 | Supplementary Fig. S7 and S9 |
| AAACGU | >20 | Supplementary Fig. S8 and S9 |

\* MD simulations of truncated NIX\_RRM-3 model.

Analysis of the NIX_RRM-3/AGACGU complex demonstrated that the RNA bases A1 and G2 (<u>AG</u>ACGU) interact mainly with Arg73 and Glu77 (Fig. 3A and C), residues that are located in the unstructured C-terminus of the protein. Evaluation of the RMSF of NIX_RRM-3 and NIX_RRM-3/AGACGU complex showed that the movement of the C-terminus is reduced significantly upon RNA binding. To investigate the interactions between the RNA and the C-terminus of the NIX_RRM-3 model the protein was truncated, the last eight C-terminal residues (PEKKVFKN) were removed, and subsequent MD simulations with AGACGU were performed. Interestingly, the RMSF of the truncated complex increased compared with the full-length NIX_RRM-3/AGACGU complex (Supplementary Fig. S7) with the RNA sequence (AGACGU) moving > 20 Å away from the starting position. These results demonstrate the importance of the protein C-terminus for the interaction with RNA. Additionally, the importance of G2 (A<u>G</u>ACGU) for the interaction with NIX_RRM-3 was investigated, thus guanine (G2) was replaced by adenine and new MD simulations were performed, the analysis indicated the formation of an unstable complex (RRM-3/AAACGU) (Table 1 and supplementary Fig. S8).

**Fig. 3.**
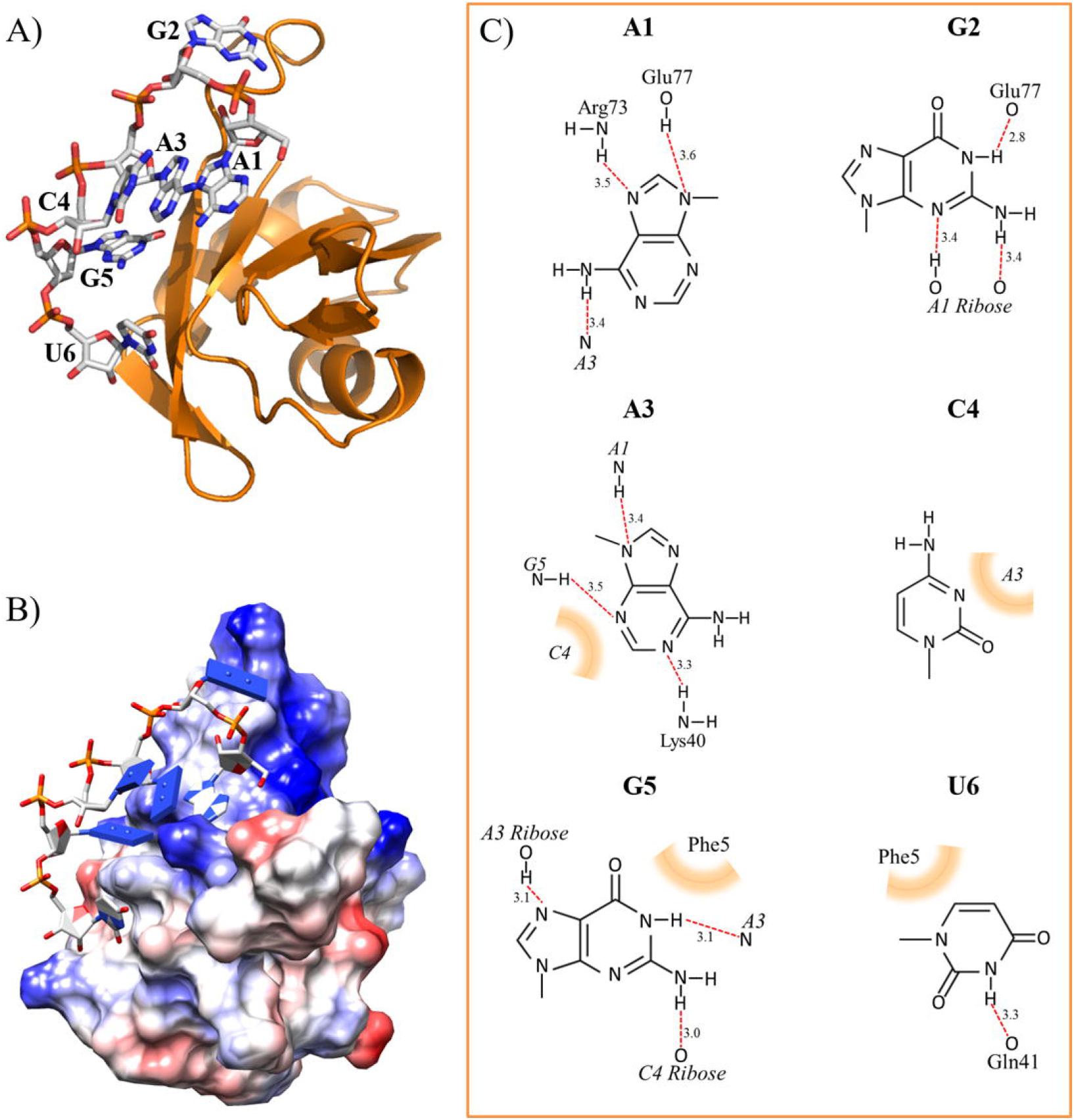
*A. aegypti* NIX_RRM-3/AGACGU complex after MD simulations. (A) NIX_RRM-3/AGACGU complex, protein in ribbon view and the RNA is presented as sticks. (B) NIX_RRM-3/AGACGU complex in a Coloumbic surface representation, positive surface in blue and negative surface in red, RNA is in sticks. (C) Interaction of AGACGU with NIX_RRM-3, single base interactions are shown. The amino acids involved in hydrogen bonds and intramolecular hydrogen bonds between the RNA bases are highlighted (red dotted lines) and hydrophobic interactions are emphasized by orange bands.

### 3.4 AGACGU interaction with *A. aegypti* NIX_RRM-3 model

The interaction of AGACGU with NIX_RRM-3 induces secondary structural changes in the protein model. The α-helix content increased from 15 to 22% and the β-strand content increased from 28 to 32 % respectively whereas the random coiled amount decreased from 40 to 29 % in the model of the NIX_RRM-3 complex compared to the unbound protein. Upon RNA binding, an additional β-strand was formed and the RNP1 containing β-3 in the free protein becomes β-4 in the protein/RNA complex (Supplementary Fig. S10A and B).

AGACGU also undergoes conformational changes during the interaction with NIX_RRM-3 (Supplementary Fig. S10C and D). Amino acid residues interacting with AGACGU are Phe5, Lys40, Gln41, Arg73 and Glu77 as indicated in Fig. 3A-C.

Interestingly, three bases in the AGACGU sequence change conformation upon binding to NIX_RRM-3. A1, G2 and G5 adopted an unusual *syn* conformation (Fig. 4); A1 is connected by hydrogen bonds with Arg73 and Glu77 and an intramolecular hydrogen bond with A3. G2 interact with Glu77 and with the sugar moiety of A1. G5 is stabilized via hydrophobic interactions with Phe5 of the protein and with intramolecular hydrogen bonds of the A3 and C4 sugar moieties (Fig. 3C).

**Fig. 4.**
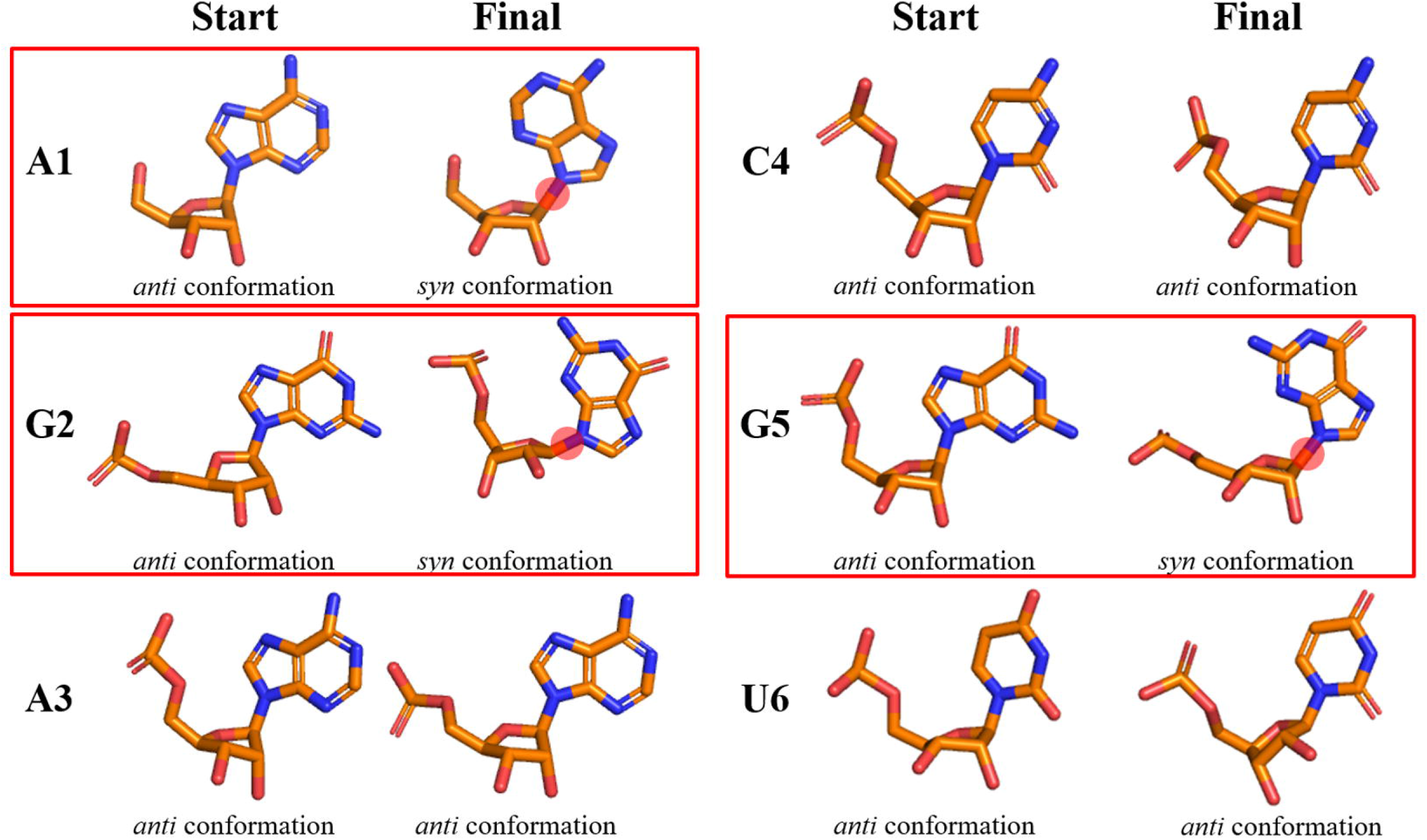
Conformational changes of the AGACGU nucleotide bases after binding to NIX_RRM-3. A1, G2 and G5 changed from *anti* to *syn* conformation. The red sphere demonstrate the rotation of the χ angle.

The *syn* conformation of RNA bases in RNA/RRM complexes occur in several examples, A4 in the 5’-AGAA-3’ sequence interacting with human TRA2B [29]; A2 of the 5’-CAUC-3’ RNA sequence bound by the Srp RRM [31]; A12 from the 5’-CAA-3’triplet interacting with RBMY RRM [32]; guanines are noted to adopt a *syn* conformation during binding with RRM proteins [33–35]. A default RNA binding sequence for an RRM that might be a dinucleotide A/C-G in N1-N2 position has been reported [28]; this is the same sequence, which we observed for the <u>AG</u>ACGU interaction with NIX_RRM-3.

Human TRA2B and PABPC1 protein structures in complex with RNA suggested that RNA mainly interacts with the core βαββαβ, formed by β-strand 1 and 3, both carrying RNP2 and RNP1 [29,30]. In this region, a typical sequence of aromatic residues in RRM proteins can be observed which are involved in the interaction with the RNA. The aromatic amino acid residues on the β1 and β4 surface Phe5, Tyr42, Phe44 and Tyr46 of NIX_RRM-3 show similarities compared with that of human TRA2B (Fig.5A and B).

**Fig. 5.**
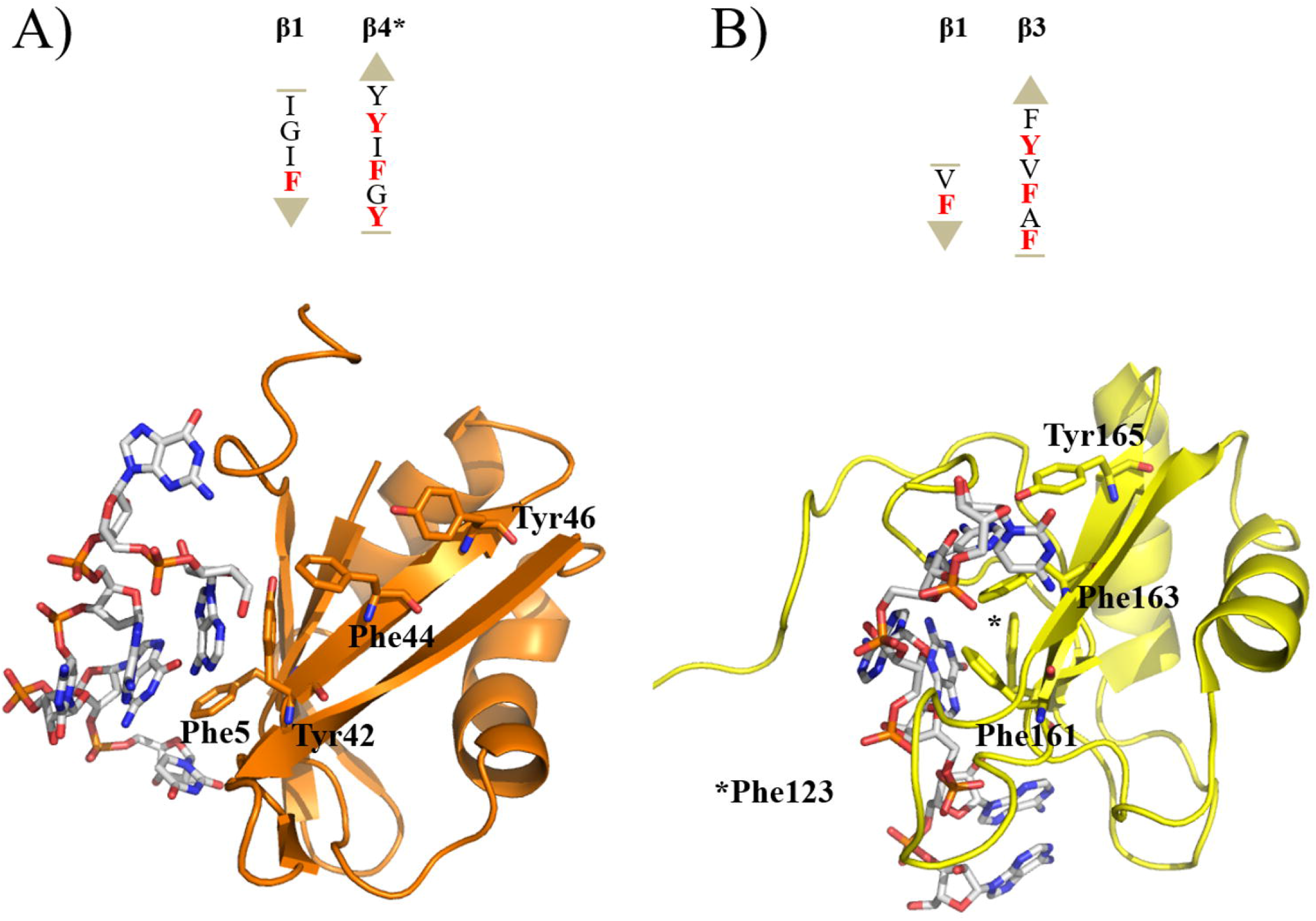
Aromatic residues in the βαββαβ core of RRM proteins that are involved in the RNA interaction. Top panel: Schematic representation of the aromatic amino acids in β-sheet 1 and β-sheet 3/4. Asterisk: After RNA interaction with NIX_RRM-3 an additional β-sheet is formed and β-3 numeration change to β-4. Down panel: tertiary structures of RRMs with RNA. (A) NIX_RRM-3. (B) Human TRA2B (PDB entry: 2KXN).

Comparison of the aromatic amino acid sequence elements of the human TRA2B in β1 and β3 with the NIX_RRM-3 model showed that Phe161 is replaced by Tyr42 (Fig. 5A and B). Tyr 42 in NIX_RRM-3 seems to have an important function in the stabilization of the RNA binding region while the residues of the C-terminus (Arg73 and Glu77) seem to interact with A1 and G2. Both residues fluctuate in the NIX_RRM-3 model (Fig. 6A).

**Fig. 6.**
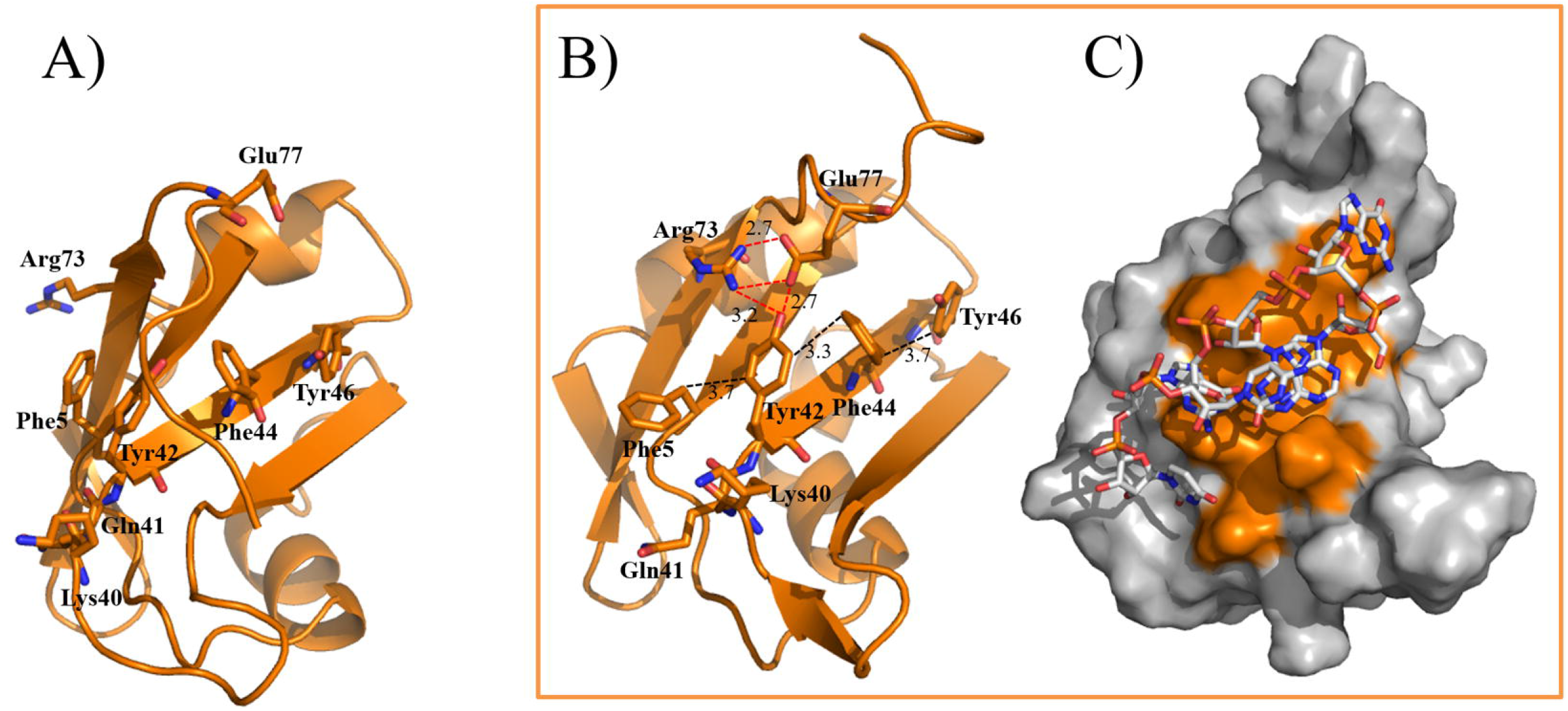
Interaction network between aromatic residues in the βαββαβ core of NIX_RRM-3 and amino acids involved in the interaction with AGACGU. (A) Tertiary structure of NIX_RRM-3. (B) Tertiary structure of NIX_RRM-3/AGACGU complex without the RNA sequence. (C) Interaction surface of NIX_RRM-3/AGACGU complex; the aromatic residues of the βαββαβ core and the amino acids involved in the interaction with AGACGU are colored (orange).

Interestingly, after interacting with AGACGU, the C-terminal residue Arg73 interacts mainly with A1 and Glu77 with both bases (A1 and G2). Furthermore, Arg73 and Glu77 form bifurcated hydrogen bonds with each other and one with Tyr42 with a distance range of 2.7 to 3.2 Å to stabilize the protein/RNA complex (Fig. 6B).

Fig. 6C demonstrates the surface of interaction of the NIX_RRM-3/AGACGU complex. It is likely that the interaction network between NIX_RRM-3 residues Tyr42, Arg73 and Glu77 forms a specific binding region for AG RNA sequences. We suggest that the C-terminus of NIX_RRM-3 is important to identify specific RNA sequences. When RRMs bind to the target RNA, N- and C-terminus of the protein may be involved in recognition and binding processes, which is typical for RRM proteins [36–38].

## 4. Conclusion

The spread of virus diseases (Dengue, Zika, Yellow fever and Chikungunya virus) transmitted by mosquitoes such as *Aedes aegypti* is an event that is occurring with greater frequency in warmer, developing countries. Low investment in fragile health systems generates a perfect setting for infectious diseases. Global climatic, environmental and ecological changes provide for the mosquitoes a favorable environment by breaking their natural and ecological boundaries and becoming established in new geographic locations. The lack of an effective antiviral treatments or vaccines for Dengue, Zika, Yellow fever and Chikungunya virus infections means that vector control is a highly attractive, alternate strategy to safeguard populations against infections. However, the efficacy of insecticide-based vector control is limited due to elevated levels of insecticide resistance in vector insect populations, lending further urgency to efforts in developing novel vector control strategies [39,40]. One promising approach, sterile insect technique (SIT), has been successfully used in past insect pest eradication programs. The new world screwworm (*Cochliomyia hominivorax*), the mediterranean fruit fly (*Ceratitis capitate*), and the tsetse fly (*Glossina morsitans*) have been successfully targeted with this technique [41–43].

There has been substantial interest in the manipulation of mosquito sex development since only female mosquitoes spread viral diseases during a blood meal. Male development in *A. aegypti* is initiated by male factors and these factors would be useful in implementing vector control strategies where developing female mosquitoes are converted into males. A new identified *A. aegypti* gene, NIX, possess typical characteristics of a male factor [14]. To date, no detailed information is available about the *A. aegypti* NIX interaction with the target RNA on the molecular level. *A. aegypti* NIX contain three RNA recognition domains and NIX_RRM-3 contains an extended C-terminal tail when compared with NIX_RRM-1 and −2. We suggest that this tail is important for the recognition of a motif in the target RNA, where the C-terminus scans the RNA until the target motif, AG, is recognized. This interaction induces conformational changes and the three RRM domains of *A. aegypti* NIX bind to the target RNA (Fig. 7).

**Fig. 7.**
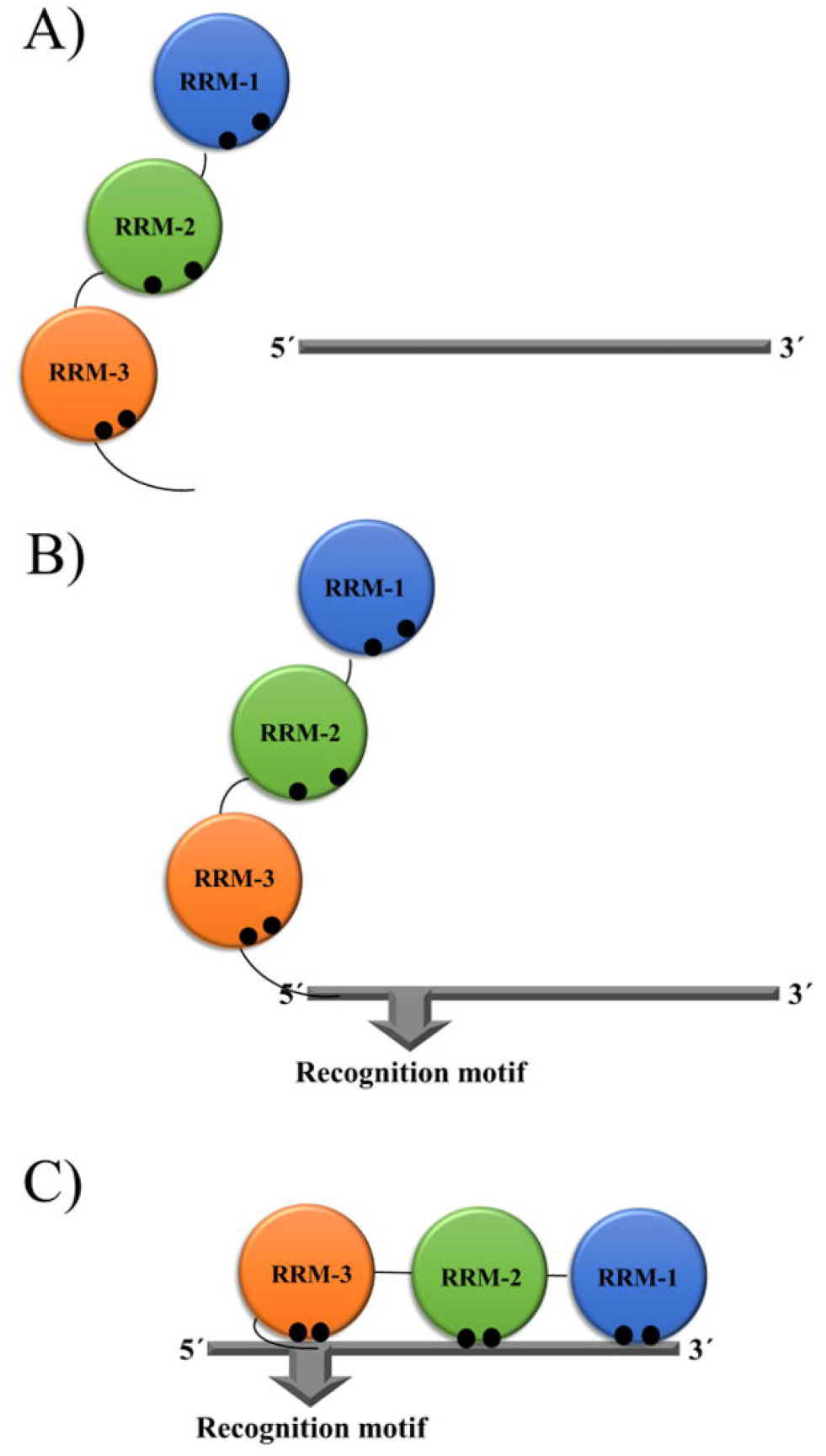
Suggested mode of recognition and interaction of *A. aegypti* NIX with target RNA. NIX_RRM1 (blue), NIX_RRM-2 (green), NIX_RRM-3 (orange), RNPs (black circles) and RNA (grey). (A) *A. aegypti* NIX and free target RNA. (B) C-terminus of NIX_RRM-3 scan the target RNA sequence for a defined recognition motif (e.g. AG). (C) After recognition of the RNA motif, the interaction of NIX_RRM-3 RNP1 and RNP2 with the target RNA can induce conformational changes in the protein and NIX_RRM-2; RRM-1 could bind to the RNA.

## Aknowledgements

We wish to thank Prof. Dr. D. Willbold, Dr. M. Schwarten, Dr. S. Feuerstein and N. Bleffert for useful discussion and for revising the manuscript. This research was supported by grants from CNPq [Grant numbers 309940/2019-2, 435913/2016-6, 401270/2014-9, 307338/2014-2], FAPESP [Grant numbers 2016/129040, 2018/07572-3, 2018/126590, 2019/05614-3], Fundect [23/200.307/2014], CAPES and PROPe UNESP.

## Author contributions

M.A.C., R.J.E and K.C.B performed the In silico analysis of NIX and NIX_RRM-3. D.S.O. and M.S.A. performed the molecular docking and MD simulations. M.A.C. and R.J.E analyzed the computational experiments. R.K.A. guided the research. M.A.C. and R.J.E coordinated the research and planned all the experiments. M.A.C. and R.J.E wrote the manuscript and all authors read and took part in revising the final version.

