## Supplementary Material for "*In silico* investigation of *Aedes aegypti* male-determining factor (NIX): RNA recognition motif-3, structural model and selective nucleic acid binding mode"

^a^Multiuser Center for Biomolecular Innovation, Departement of Physics, Instituto de Biociências
Letras e Ciências Exatas (Ibilce), Universidade Estadual Paulista (UNESP), São Jose do Rio Preto-SP, 15054-000, Brazil.

^b^Institute of Biological Information Processing (IBI-7: Structural Biochemistry), Forschungszentrum Jülich, Jülich, Germany

^c^Federal University of Tocantins, Araguaína-TO, Brazil.

^d^Institute of Physics, Federal University of Mato Grosso do Sul, Campo Grande-MS, Brazil.

* Both authors contributed equally

Table of contents

Figure S1. Time dependent modifications of *A. aegypti* Nix_RRM-3 during MD simulations.

Figure S2. Structural comparison of *A. aegypti* Nix_RRM-3, human TRA2B and PABPC1.

Figure S3. Homology models of *A. aegypti* NIX_RRM 1-3 domains.

Figure S4. RNA binding of human PABPC1 and TRA2B.

Figure S5. Time dependent modifications of *A. aegypti* Nix_RRM-3 AAGAAC complex during MD simulations.

Figure S6. Time dependent modifications of *A. aegypti* Nix_RRM-3 AGACGU complex during MD simulations.

Figure S7. Time dependent modifications of truncated *A. aegypti* Nix_RRM-3 AGACGU complex during MD simulations.

Figure S8. Time dependent modifications of *A. aegypti* Nix_RRM-3 AAACGU complex during MD simulations.

Figure S9. Starting and final positions of RNA molecules before and after MD simulations with *A. aegypti* Nix_RRM-3.

Figure S10. Secondary structure changes of *A. aegypti* Nix_RRM-3 and AGACGU after MD simulations.

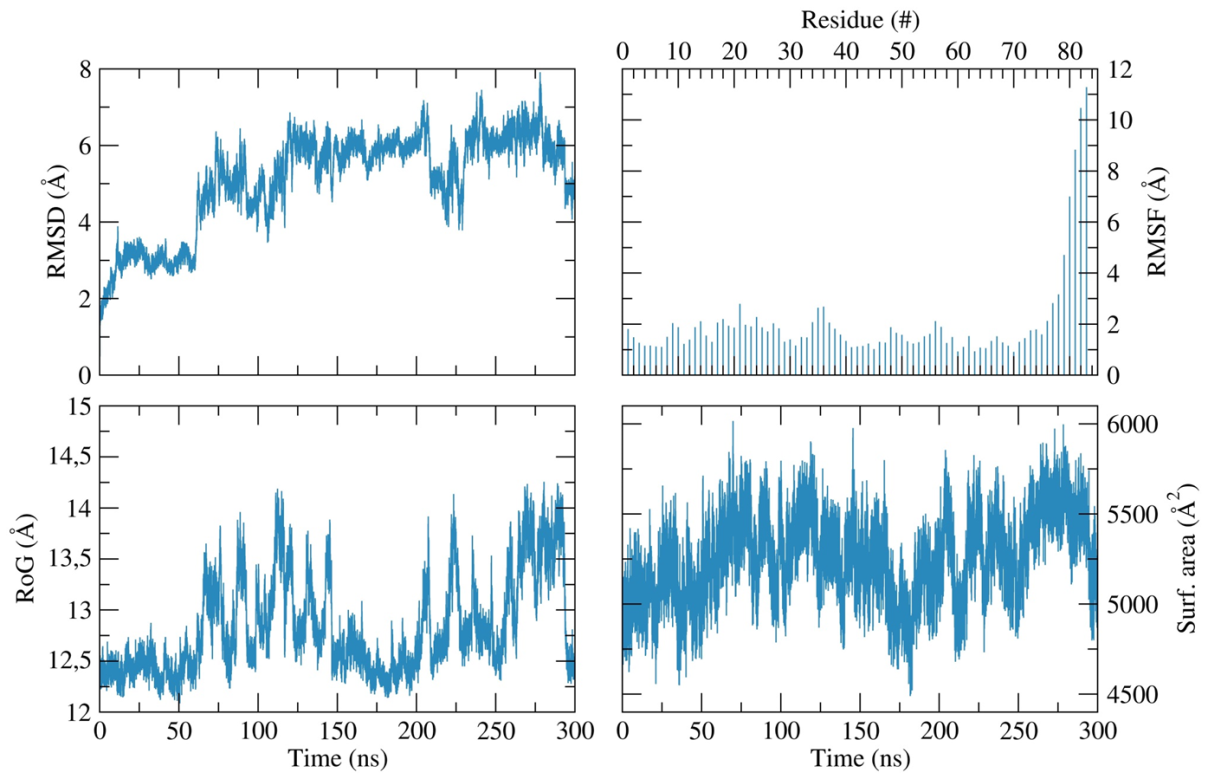

**Fig. S1.** Time dependent modifications of the Nix_RRM-3 model during MD simulations; RMSD as function of time.

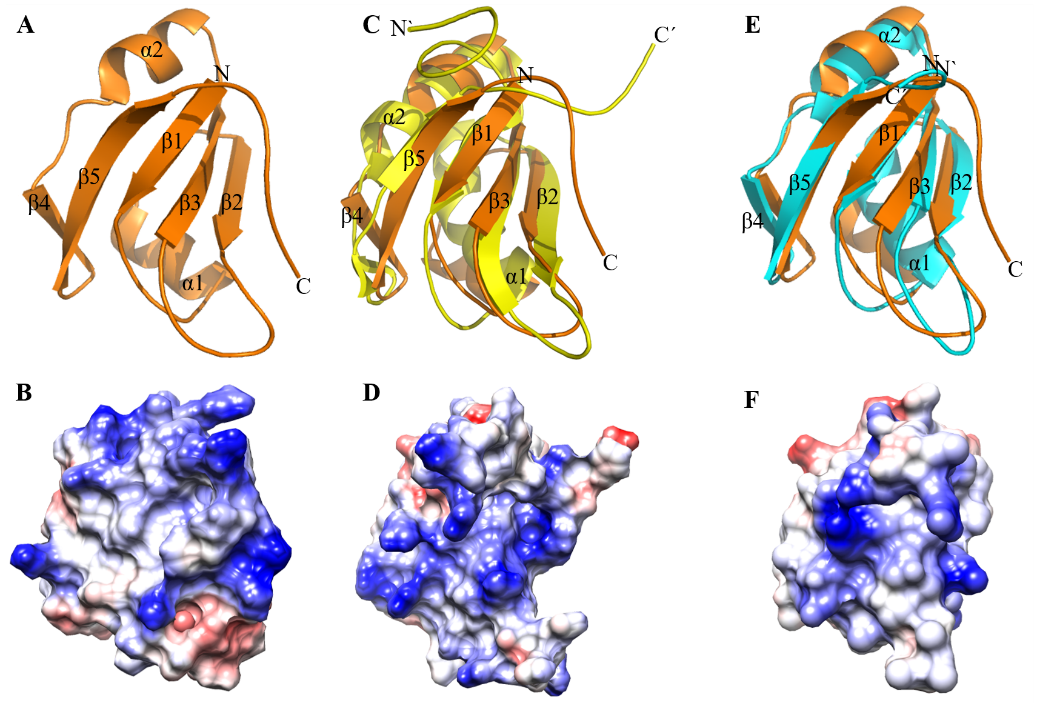

**Fig. S2.** Structural comparison of *A. aegypti* Nix_RRM-3 (orange), human TRA2B (yellow) and PABPC1 (turquoise); Ribbon and coulombic surface view. Positive surface in blue and negative surface in red. (A and B) *A. aegypti* Nix_RRM-3.(C and D) Human TRA2B (PDB entry: 2KXN). (E and F) Human PABPC1 (PDB entry: 1CVJ).

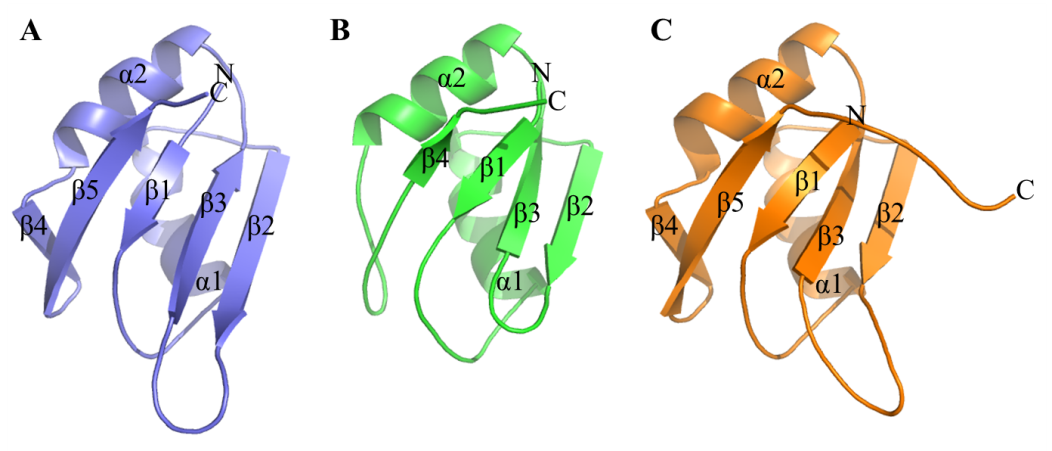

**Fig. S3.** Homology models of *A. aegypti* NIX_RRM 1-3. (A) NIX_RRM-1. (B) NIX_RRM-2. (C) NIX_RRM-3.

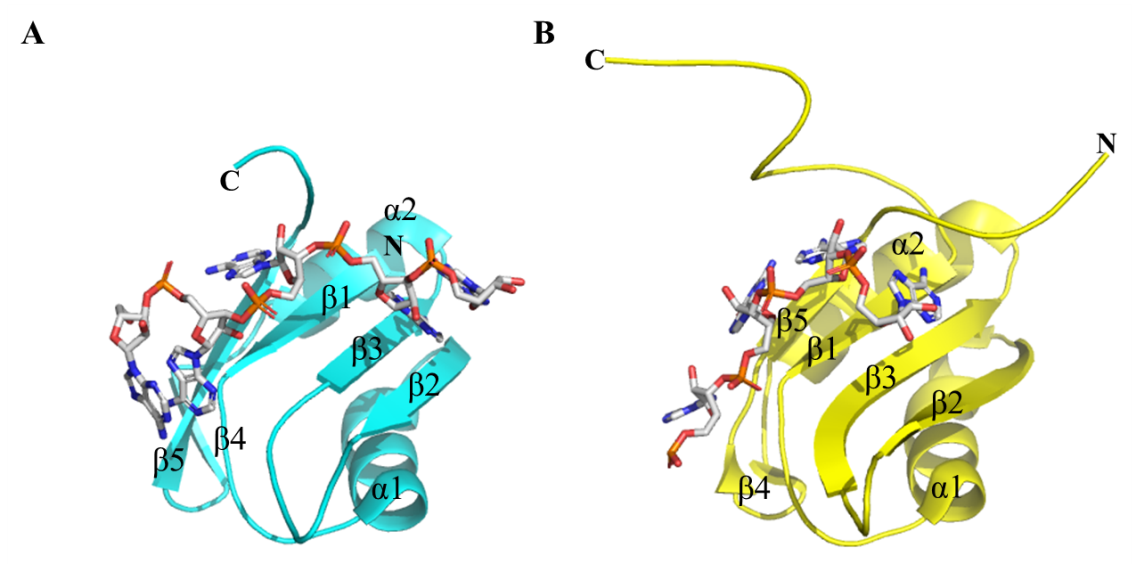

**Fig. S4.** RNA binding of human PABPC1 (turquoise) and TRA2B (yellow). (A) Human PABPC1 (PDB entry: 1CVJ). (B) Human TRA2B (PDB entry: 2KXN).

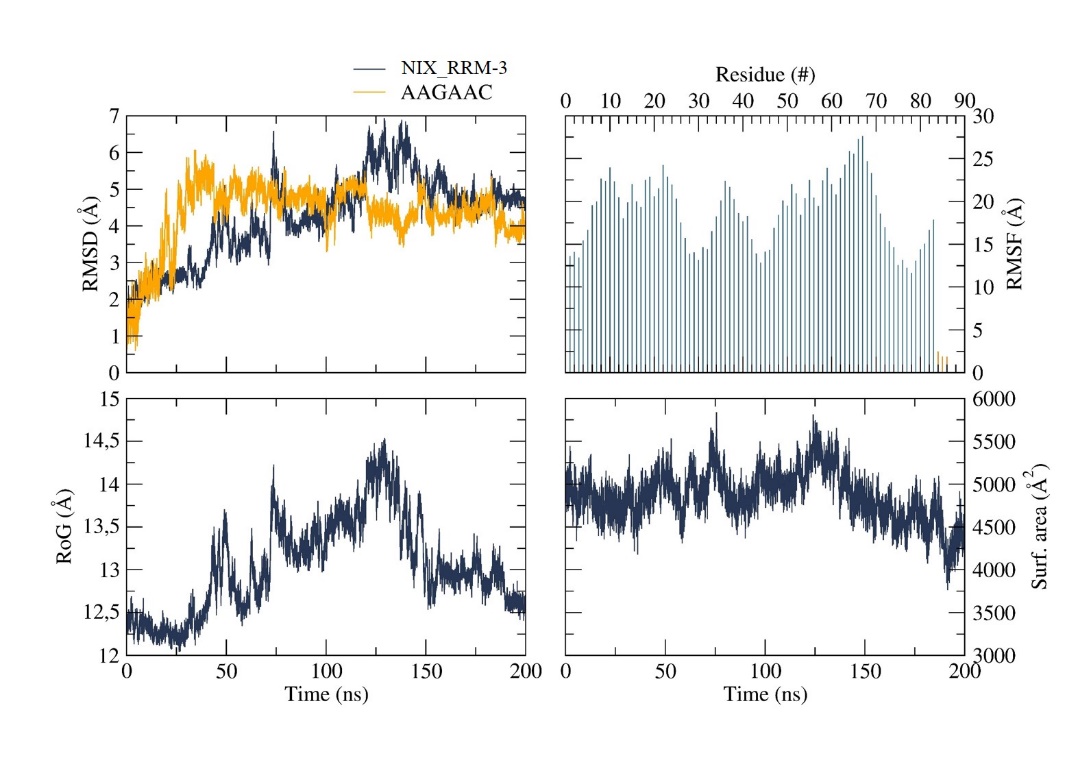

**Fig. S5.** Time dependent modifications of *A. aegypti* Nix_RRM-3/AAGAAC complex during MD simulations Nix_RRM-3 (dark blue) and AAGAAC (yellow); RMSD as function of time.

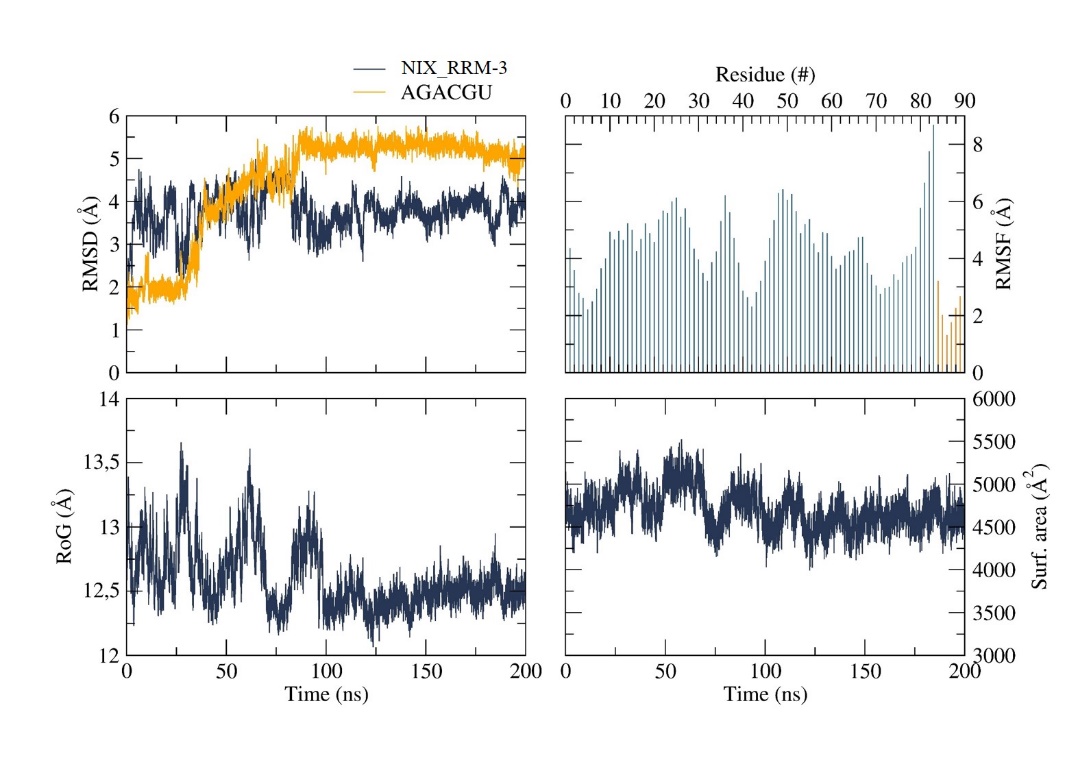

**Fig. S6.** Time dependent modifications of *A. aegypti* Nix_RRM-3/AGACGU complex during MD simulations Nix_RRM-3 (dark blue) and AGACGU (yellow); RMSD as function of time.

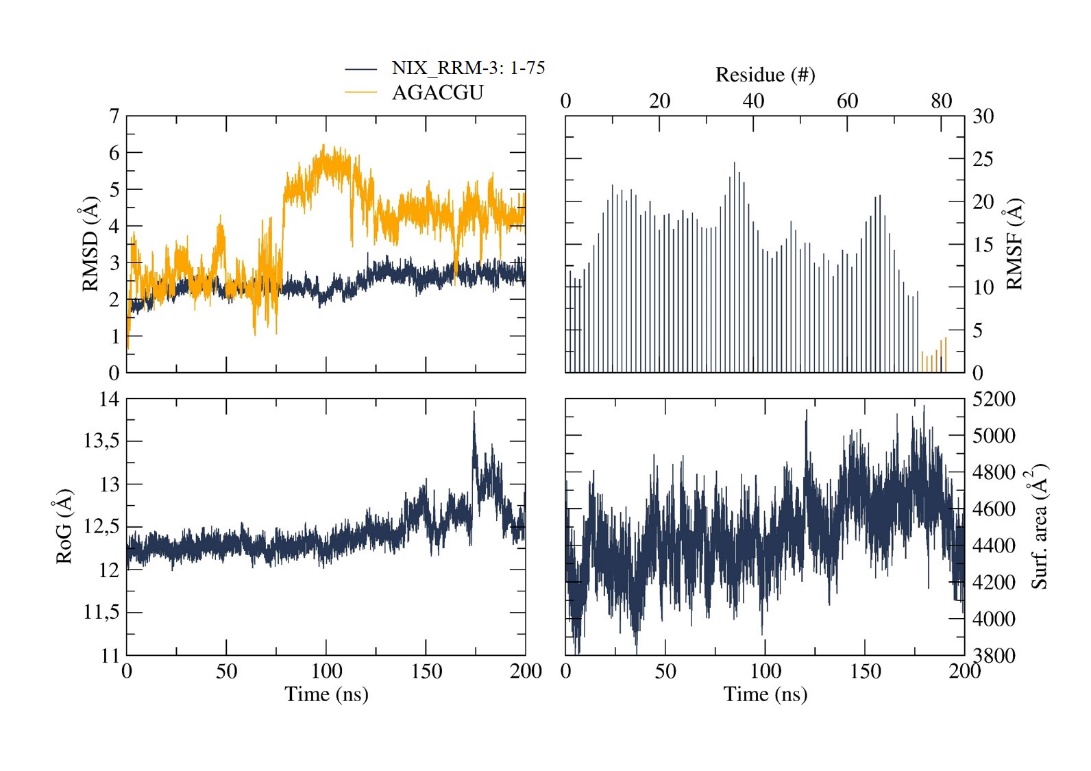

**Fig. S7.** Time dependent modifications of truncated *A. aegypti* Nix_RRM-3/AGACGU complex during MD simulations truncated Nix_RRM-3 (dark blue) and AGACGU (yellow); RMSD as function of time.

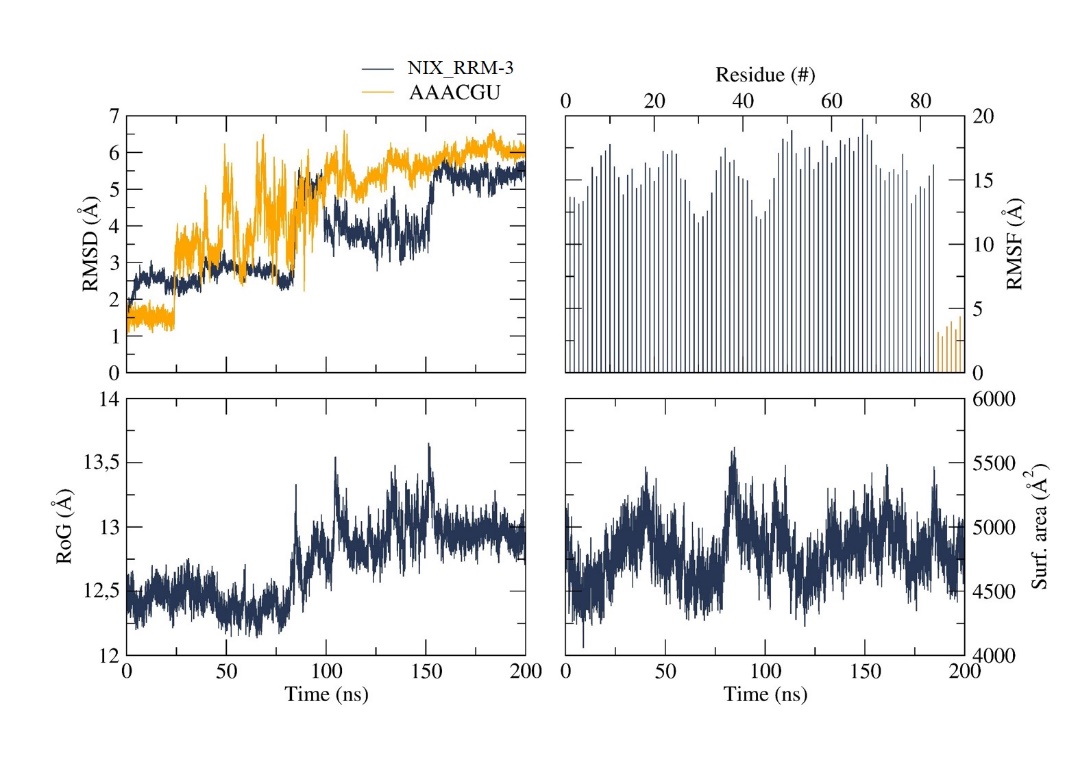

**Fig. S8.** Time dependent modifications of *A. aegypti* Nix_RRM-3/AAACGU complex during MD simulations Nix_RRM-3 (dark blue) and AAACGU (yellow); RMSD as function of time.

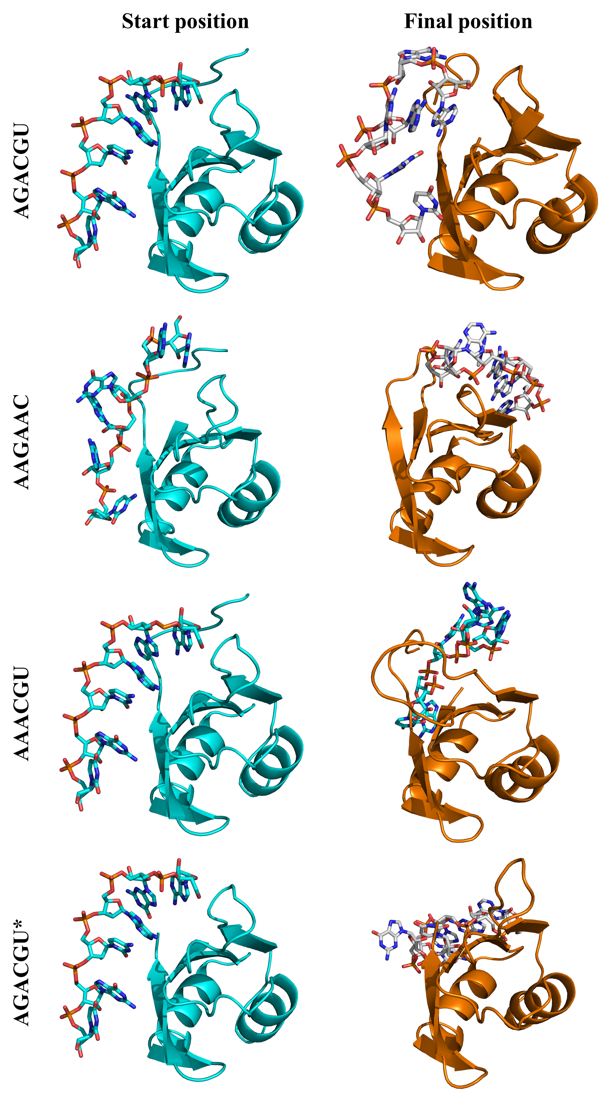

**Fig. S9.** Starting and final positions of RNA molecules before and after MD simulations with *A. aegypti* Nix_RRM-3 model. Left panel: Nix_RRM-3 model (turquoise) with RNA start position for MD simulations. Right panel: Nix_RRM-3 model (orange) with RNA final position after MD simulations. Asterisks label the truncated Nix_RRM-3 model.

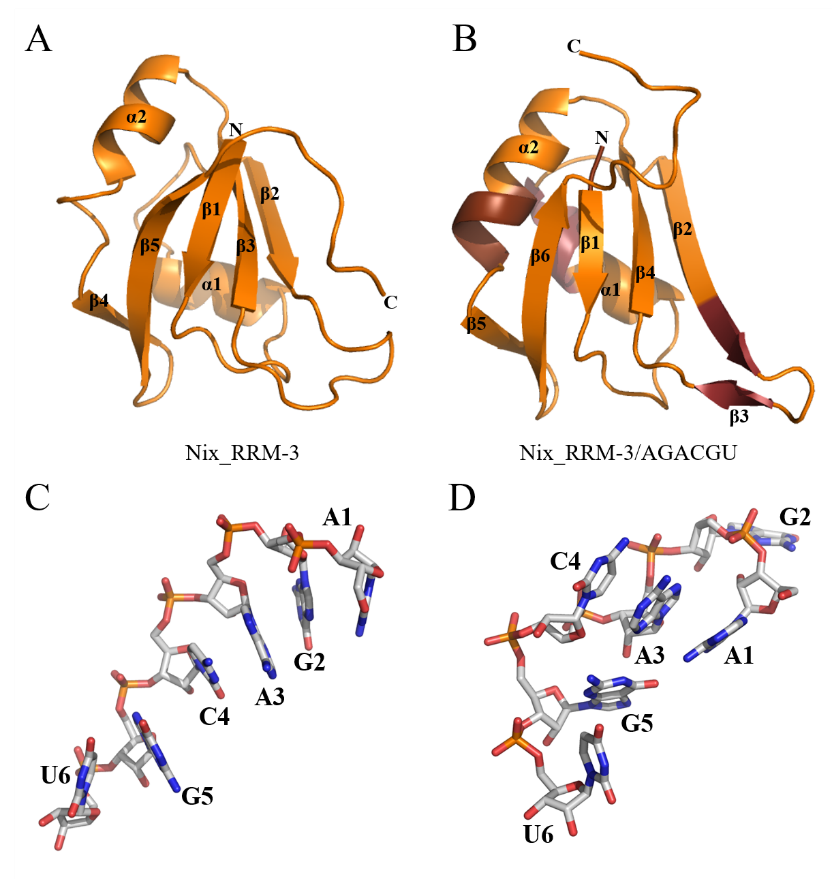

**Fig. S10.** Secondary structure changes of *A. aegypti* Nix_RRM-3 and AGACGU after MD simulations. Protein in ribbon view and DNA is shown in sticks. (A) Nix_RRM-3 starting model. (B) Nix_RRM-3 after interaction with AGACGU (RNA not shown), protein regions undergoing secondary structure changes are colored brown. (C) AGACGU starting structure. (D) AGACGU final structure after interaction with Nix_RRM-3.
